# A simple permutation-based test of intermodal correspondence

**DOI:** 10.1101/2020.09.10.285049

**Authors:** Sarah M. Weinstein, Simon N. Vandekar, Azeez Adebimpe, Tinashe M. Tapera, Timothy Robert-Fitzgerald, Ruben C. Gur, Raquel E. Gur, Armin Raznahan, Theodore D. Satterthwaite, Aaron F. Alexander-Bloch, Russell T. Shinohara

## Abstract

Many key findings in neuroimaging studies involve similarities between brain maps, but statistical methods used to measure these findings have varied. Current state-of-the-art methods involve comparing observed group-level brain maps (after averaging intensities at each image location across multiple subjects) against spatial null models of these group-level maps. However, these methods typically make strong and potentially unrealistic statistical assumptions, such as covariance stationarity. To address these issues, in this paper we propose using subject-level data and a classical permutation testing framework to test and assess similarities between brain maps. Our method is comparable to traditional permutation tests in that it involves randomly permuting subjects to generate a null distribution of intermodal correspondence statistics, which we compare to an observed statistic to estimate a *p*-value. We apply and compare our method in simulated and real neuroimaging data from the Philadelphia Neurodevelopmental Cohort. We show that our method performs well for detecting relationships between modalities known to be strongly related (cortical thickness and sulcal depth), and it is conservative when an association would not be expected (cortical thickness and activation on the *n*-back working memory task). Notably, our method is the most flexible and reliable for localizing intermodal relationships within subregions of the brain and allows for generalizable statistical inference.

## 1 Introduction

Neuroimaging studies often seek to understand similarities across modalities. However, methods underlying these studies’ findings continue to vary. Naive approaches to the so-called “correspondence problem” have included simply visualizing two maps next to each other and observing whether the two maps look similar. Another approach involves the use of parametric *p*-values; for example, fitting a linear regression or estimating the correlation across locations of the cortex and using a parametric null distribution (e.g., Student’s *t*-distribution), ignoring inherent spatial autocorrelation. A third approach has been to use a partial correlation coefficient controlling for anatomical distance (Honey et al. 2009; Horn et al. 2014).

Methods that ignore spatial autocorrelation have been shown to inflate type I error rates (Markello and Misic 2021); thus, recent methodological advancements that do account for spatial correlation have been widely adopted by the neuroimaging community. A spatial permutation framework was first introduced by Alexander-Bloch et al. (2013), using spatially-constrained randomization to generate null models for testing intermodal developmental synchrony (Alexander-Bloch et al. 2013). Vandekar et al. (2015) then used spherical rotations to obtain null models of spatial alignment in testing correspondence between various cortical measurements in the human brain. The spin test was formally introduced by Alexander-Bloch et al. (2018), who used random spherical rotations of group-level or population-averaged surfaces to generate null models of spatial alignment. A surface-based permutation null model was also developed independently in the context of validating functional magnetic resonance imaging parcellations (Gordon et al. 2016; Gordon et al. 2017a; Gordon et al. 2017b).

Since the advent of the spin test, numerous neuroimaging experiments have adopted it to corroborate claims about intermodal similarities (Lefèvre et al. 2018; Paquola et al. 2019; Schmitt et al. 2019; Shafiei et al. 2020; Cui et al. 2020; Stoecklein et al. 2020) and have extended its implementation to regionally parcellated maps (Váša et al.2018; Vázquez-Rodríguez et al. 2019; Baum et al. 2020; Cornblath et al. 2020). The spin test was certainly an improvement compared with prior approaches; however, deriving frequentist properties of inference across datasets using this method depends on strong and often unrealistic statistical assumptions, such as stationarity of the covariance structure. Other critiques of the spin test include its limited use in the context of cortical surface maps without a straightforward extension to volumetric data, its lack of flexibility for incorporating subject-level data by accommodating only two (group-level) maps, and that it cannot be used to test correspondence in small regions of the brain. Another limitation of the formal support for the spin test presented in Alexander-Bloch et al. (2018) is its ad hoc treatment of the medial wall, which is an artifact of generating a topologically spherical surface that does not represent neuroanatomical features (Dale et al. 1999).

More recently, Burt et al. (2020) proposed Brain Surrogate Maps with Autocorrelated Spatial Heterogeneity (BrainSMASH), using generative modeling to obtain null models seeking to preserve the spatial autocorrelation structure of the observed data. Burt et al.’s approach greatly improves on several aspects of the spin test in its flexibility to incorporate volumetric data, its inclusion of the medial wall, and its ability to test correspondence in small regions of the brain. Still, BrainSMASH has some limitations. When applied to subregions of the brain, one can only account for spatial autocorrelation among locations within each subregion. For instance, when testing intermodal correspondence between two maps of the left hemisphere, spatial autocorrelation among locations within that hemisphere are modeled, but correlation with locations in the right hemisphere are not. Furthermore, BrainSMASH models spatial, but not functional, relationships between locations throughout the brain.

The spin test, BrainSMASH, and other related methods recently described by Markello and Misic (2021) involve testing null hypotheses of intermodal correspondence conditional on observed group-level data (Burt et al. 2020; Alexander-Bloch et al. 2018; Vázquez-Rodríguez et al. 2019; Baum et al. 2020; Cornblath et al. 2020; Váša et al. 2018; Burt et al. 2018; Wael et al. 2020). As illustrated in Figure 1, averaging images across subjects precludes evaluation of inter-individual heterogeneities. Importantly, these previous methods may not be well-suited to account for dataset-to-dataset variability without strong assumptions about stationarity of the covariance structure. Burt et al. (2020) use a variogram-matching model that assumes stationarity and normality—assumptions that have been subject to criticism in the context of neuroimaging studies (Ye et al. 2015; Eklund et al. 2016). While these methods are currently the most sophisticated in the field, their underlying assumptions may give rise to inferential missteps, with high false positive rates and substantial variability in results (Markello and Misic 2021). Thus, careful consideration of the plausibility of these assumptions is warranted, and the appeal of a method without strong assumptions motivates our current work.

**Figure 1:**
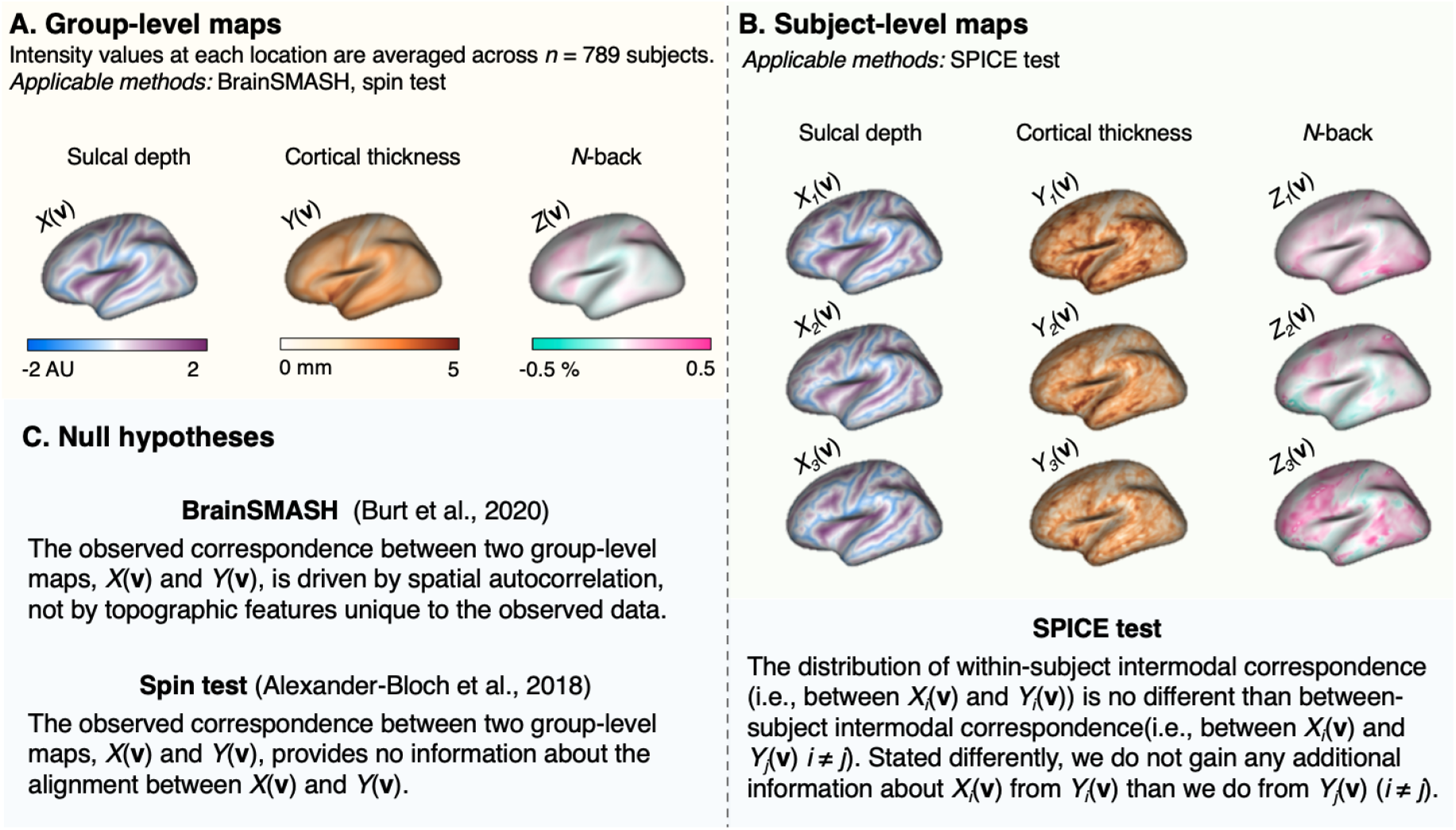
Example left hemisphere sulcal depth, cortical thickness, and *n*-back maps used for (A) the spin test and BrainSMASH and (B) the SPICE test. Brain map visualizations were generated using the R package fsbrain (Schäfer 2020b). (C) Null hypotheses for previous methods and our proposed method for testing intermodal correspondence. Of the three methods, the SPICE test is the only one to incorporate subject-level data.

In this paper, we propose the simple permutation-based intermodal correspondence (SPICE) test, a novel procedure that is intuitive, easy to implement, and does not depend on strong statistical assumptions. This test considers a setting with subjects *i* = 1,…,n for whom we observe two imaging modalities each: *X_i_*(**v**) and *Y_i_*(**v**) at locations (e.g., pixels, voxels, or vertices) indexed by **v** = [1,…,*V*]. Our null hypothesis states that the distribution of within-subject intermodal similarity between *X_i_*(**v**) and *Y*_i_(**v**) (i.e., two modalities belonging to the same subject) is no different than the distribution of between-subject intermodal correspondence—for example, between *X_i_*(**v**) and *Y_j_*(**v**) (*i* ≠ *j*). In prior studies, the Pearson correlation has been widely used to measure intermodal correspondence (Alexander-Bloch et al. 2018; Burt et al. 2020). While in this paper we adopt this commonly used statistic, in the Discussion we propose alternative measures that we hope to explore in future work.

The intuition behind the SPICE test is that, if there is a genuine anatomical correspondence between two modalities, this correspondence should be greater in brain maps derived from the same individual than from different individuals. To test the SPICE null hypothesis, we leverage subject-level data and do not require altering the spatial structure or alignment of the observed data, in contrast to BrainSMASH and the spin test. Since we will compare our proposed method to BrainSMASH and the spin test, it is important to highlight that their underlying null hypotheses are fundamentally different, though they may be used to answer related—but nevertheless different—questions about intermodal correspondence. Figure 1 outlines these three null hypotheses alongside the different types of data required for each test. We will reference the three null hypotheses throughout the paper as *H*_0(*spice*)_, *H*_0(*b-smash*)_, and *H*_0(*spin*)_.

## 2 Methods

### 2.1 The SPICE test

Formally, we express the SPICE test’s null hypothesis as:

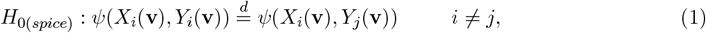

where *ψ*() represents a measure of intermodal similarity between *X* and *Y* across all locations indexed by **v**, either for the same subject or for different subjects (when *i* ≠ *j*). Adopting the Pearson correlation to measure intermodal correspondence (which has been widely used in previous work), the test of equality of means (1) is a test of whether within-subject intermodal correlation is higher than the between-subject intermodal correlation.

To test this null hypothesis, we first obtain the following empirical estimate of the mean of the left hand side of (1):

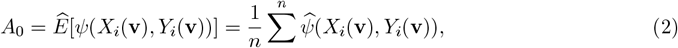

where, for the *i*th subject, the Pearson correlation across all *V* locations in the image is:

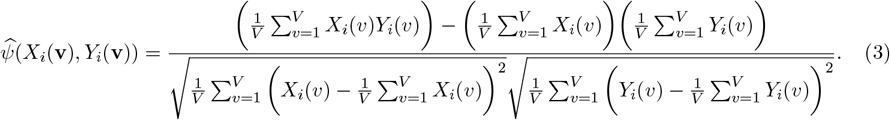

After estimating the mean within-subject intermodal correspondence (*A*_0_), we generate a null distribution for *A*_0_ by randomly shuffling the *Y* maps across subjects. Then, for the *k*th permuted sets of map pairs (*k* = 1,…,*K* permutations), we re-calculate (2) as *A_k_*, providing draws from an empirical null distribution of *A*_0_. These random permutations are illustrated in Figure 2.

**Figure 2:**
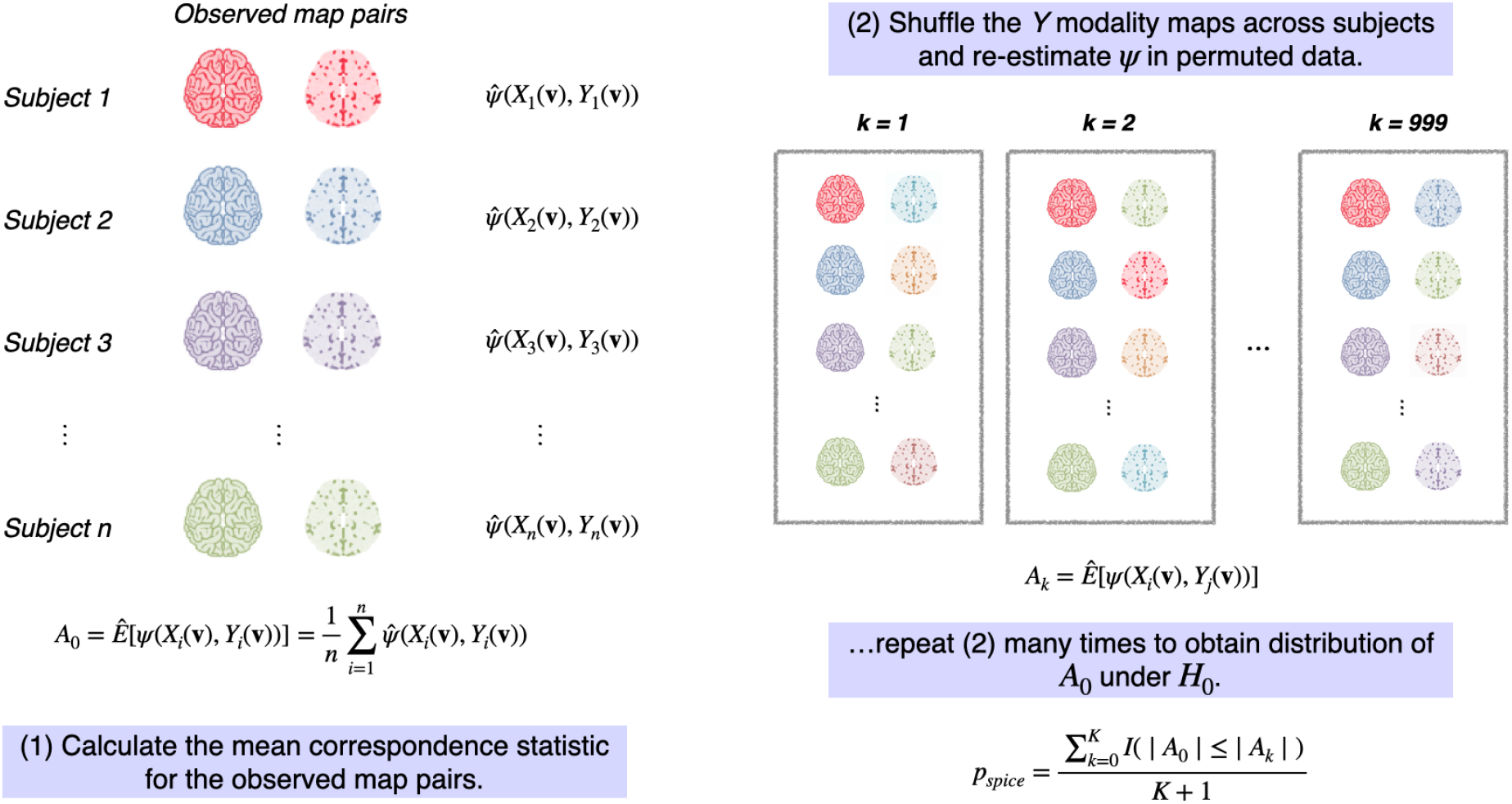
Illustration of simple permutation-based intermodal correspondence (SPICE) testing procedure. A null distribution for *A*_0_, the average within-subject correspondence statistic, is calculated by randomly permuting the *Y* maps *K* = 999 times and re-estimating *A_k_* in each permuted version of the data. A *p*-value is estimated as in (4).

A *p*-value is estimated by counting the number of permutations for which the magnitude of *A_k_* exceeds that of *A*_0_:

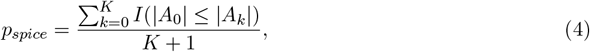

where we add 1 to the denominator to account for the identity permutation (Phipson and Smyth 2010). We reject the null hypothesis for *p_spice_* < *α*, the nominal type I error rate. Under the null hypothesis, *p_spice_* is a random variable distributed as Uniform(0,1), and we would reject *H*_0(*spice*)_ in only approximately 5% of cases. Under the alternative hypothesis, we expect *A_k_* to rarely be greater in magnitude than *A*_0_ and would reject *H*_0(*spice*)_.

### 2.2 Simulations and application in the Philadelphia Neurodevelopmental Cohort

We assess the performance of our proposed method using neuroimaging data from the Philadelphia Neurodevelopmental Cohort (PNC). The PNC is a research initiative at the University of Pennsylvania consisting of 9498 subjects between the ages of 8 and 21 who received care at the Children’s Hospital of Philadelphia and consented (or received parental consent) to participate. A subset of 1601 subjects were included in multimodal neuroimaging studies and were scanned in the same scanner at the Hospital of the University of Pennsylvania, per protocols described briefly below and in Appendix A1.

For the current study, we first consider correspondence between cortical thickness and sulcal depth—a pair of imaging measurements for which we expect to reject the null hypothesis due to well-established interdependence in structure throughout multiple brain regions shown in research from the last two decades (Vandekar et al. 2016; Vandekar et al. 2015; Sowell et al. 2004; Shaw et al. 2008), as well as in much earlier studies of neuroanatomy from nearly a century ago (Von Economo 1929). Second, we consider cortical thickness and *n*-back, a measure of working memory function (described below), to demonstrate how our method performs in a situation where intermodal correspondence would typically not be expected.

In this study, we include a subset of *n* = 789 subjects with all three image types (cortical thickness, sulcal depth, and *n*-back) meeting quality control criteria. This subset also excluded individuals with existing health conditions, who were taking psychoactive medications, had a history of psychiatric hospitalization, or other abnormalities that could impact brain structure or function.

#### 2.2.1 Acquisition and processing of multimodal imaging data

Relevant protocols for the acquisition and pre-processing of both structural and functional magnetic resonance imaging (MRI) data are described in Appendix A1. In brief, all subjects underwent MRI scans in the same Siemens TIM Trio 3 tesla scanner with a 32-channel head coil and the same imaging sequences and parameters (Satterthwaite et al. 2014; Satterthwaite et al. 2016). Cortical reconstruction of T1-weighted structural images was implemented using FreeSurfer (version 5.3). We also consider the fractal *n*-back task sequence acquired during functional MRI (fMRI) acquisition sequences, which involved presenting each subject with a series of stimuli (geometric pictures on a screen) with instructions to press a button if the current stimulus matches the *n*th previous one. In our present analysis we consider maps that represent the percentage change in activation between the 2-back and 0-back sequences, which has been previously shown to isolate executive network function (Ragland et al. 2002; Satterthwaite et al. 2013). All measurements were resampled to the fsaverage5 atlas, consisting of *V* = 10,242 vertices in each hemisphere for every subject.

#### 2.2.2 Simulations

To evaluate the performance of the SPICE test under the null and alternative hypotheses, we conduct several simulation studies. Using average cortical thickness, sulcal depth, and n-back maps from the PNC, we simulate subject-level multimodal imaging data by adding and multiplying random noise and signal to mean maps, as shown in Figure 3. While we generate three hypothetical imaging modalities per simulated subject, to test the null hypothesis of intermodal correspondence we consider only two such modalities at a time (hence Figure 3 illustrates the simulation set-up when mean cortical thickness and mean sulcal depth maps are used). The null or alternative hypothesis is true depending on 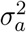, the variance of the simulated signal (*a_i_* term in Figure 3), and the null hypothesis is more difficult to reject for larger values of 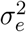, the variance of the simulated noise (elements of *E*_*i*1_(**v**) or *E*_*i*2_(**v**) in Figure 3).

**Figure 3:**
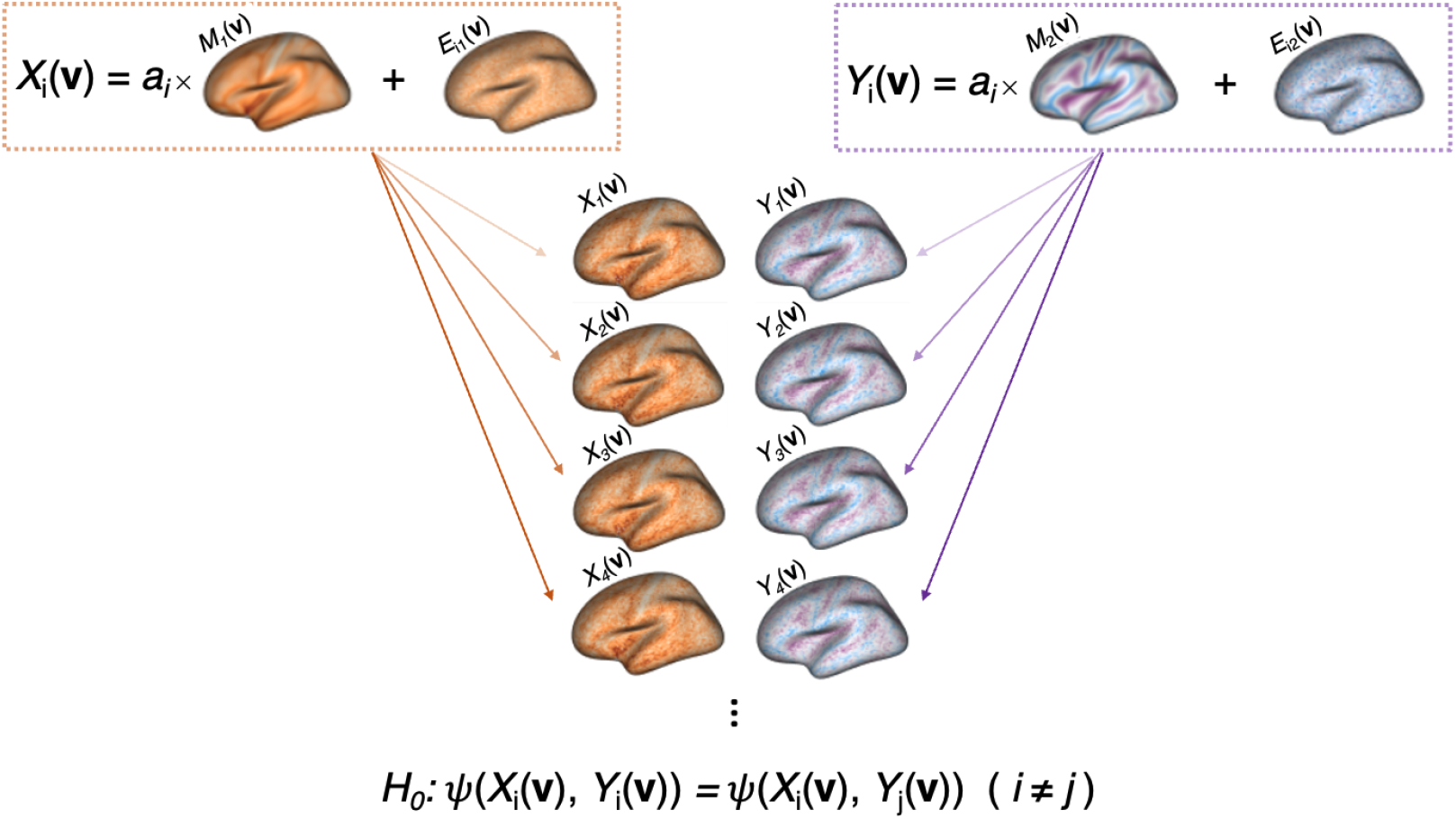
Illustration of bi-modal image simulation. Subject-level images are derived from average cortical thickness (*M*_1_(**v**)) and sulcal depth (*M*_2_(**v**)) data from the Philadelphia Neurodevelopmental Cohort. For illustrative purposes, the example below shows *M*_2_(**v**) as mean sulcal depth, but we also consider a simulation setting using mean *n*-back as the second population-level map. Subject-level images for *i* = 1,…,*n* subjects are simulated as *X*_i_(**v**) = *a_i_* × *M*_1_(**v**) + *E*_*i*_(1)__(**v**) and *Y_i_*(**v**) = *a_i_* × *M*_2_(**v**) + *E*_*i*(2)_(**v**), where *a_i_* and the elements of *E*__*i*(1)__(**v**) and *E*_*i*(2)_(**v**) are normally distributed, with mean and variance parameters specified in Section 2.2.2. The null hypothesis is true when the variance of *a_i_*, 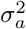 is equal to 0. Otherwise, we expect to reject *H*_0_, with test power varying according to other parameter values.

For the *i*th subject, we simulate *X_i_*(**v**) and *Y_i_*(**v**), which have a one-to-one correspondence at all locations in the fsaverage5 atlas space, indexed by **v** = [1,…, 10,242]. We define *X_i_*(**v**) = *a_i_* × *M*_1_(**v**) + *E_i_*(_1_)(**v**) and *Y_i_*(**v**) = *a_i_* × *M*_2_(**v**) + *E*_*i*(2)_(**v**), where *M*_1_(**v**) is the mean cortical thickness map, *M*_2_(**v**) is either the mean sulcal depth map (in which Cor(*M*_1_(**v**), *M*_2_(**v**)) = −0.15) or the mean *n*-back map (Cor(*M*_1_(**v**), *M*_2_(**v**)) = −0.04). We consider two sets of simulations, where *M*_1_(**v**) is mean cortical thickness in both cases, and *M*_2_(**v**) is mean sulcal depth in one setting and mean *n*-back in the other. The purpose of these two separate settings is to consider the degree to which population-level modality similarity (e.g., correlation between *M*_1_(**v**) and *M*_2_(**v**)) impacts the power of our test.

Since *X_i_*(**v**) and *Y_i_*(**v**) share *a_i_* (distributed 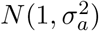), the SPICE test’s null hypothesis (1) is true when the variance of *a_i_*, 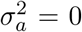 because in this case, the covariation between *X_i_*(**v**) and *Y_i_*(**v**) is no stronger than it is between *X_i_*(**v**) and *Y_j_*(**v**) (*i* ≠ *j*). The *V*-dimensional surfaces *E*_*i*(1)_(**v**) and *E*_*i*(2)_(**v**) consist of elements that are independent and identically distributed as 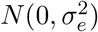.

We consider sample sizes of *n* = 25, 50, and 100, 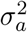 ranging from 0 to 3.0, and 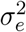 of 0.5, 1.5, 3.0, or 6.0. While the signal-to-noise ratio varies with the magnitude of intensities at different locations in *M*_1_(**v**) and *M*_2_(**v**) (i.e., higher magnitudes of intensities at each location would multiplicatively increase the variance of the signal at those locations), the relative size of 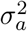 and 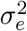 translates to varying degrees of statistical power when we conduct tests of intermodal correspondence between simulated surfaces. For instance, higher variance of the signal 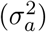 and lower variance of the noise 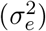 would produce more signal than noise, adding to the similarity between *X_i_*(**v**) and *Y_i_*(**v**) (from the same subject). In contrast, lower 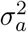 relative to 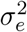 would mute the dissimilarity between *X_i_*(**v**) and *Y_j_*(**v**) (*i* ≠ *j*) (more noise than signal) and induce lower statistical power.

When 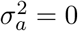, we repeat 5000 simulations of each combination of *n*, 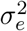, and *M*_1_(**v**) and *M*_2_(**v**) to estimate type I error as the proportion of simulations for which *p_*spice*_* < 0.05. In addition, from 5000 simulations of each combination of parameters where 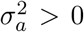, we estimate power by the proportion of simulations for which *p_spice_* < 0.05. R code for implementing the SPICE test and reproducing our simulations may be found at https://github.com/smweinst/spice_test.

#### 2.2.3 Comparing the SPICE test with previous methods

In assessing intermodal correspondence in the PNC, we compare the performance of our method to that of the spin test and BrainSMASH. Since both the spin test and BrainSMASH require group-level surfaces and their underlying null hypotheses differ, our goal here is to assess whether the general conclusions of their respective null hypotheses (Figure 1) agree with one another. Given the developmental focus of the dataset and prior research suggesting that intermodal coupling may change throughout development (Vandekar et al. 2016), we apply each method in imaging data within seven age subgroups (group-averaged for the spin test and BrainSMASH, and subject-level for the SPICE test). We also apply each method to test for intermodal correspondence in the full group of *n*=789 subjects. We account for multiple comparisons by using a Bonferroni-adjusted *p*-value threshold for statistical significance (0.05/16 = 0.003) to correct for the seven age groups and one group including all subjects, times two hemispheres tested within each group.

The spin test is implemented using publicly available MATLAB code (https://github.com/spin-test). We generate 1000 null models of spatial alignment through random rotations of cortical thickness maps within each hemisphere, after removing vertices identified to be part of the medial wall. The same “spun” maps are used to test *H*_0(*spice*_) of cortical thickness versus sulcal depth as well as cortical thickness versus *n*-back.

BrainSMASH is implemented in Python (https://github.com/murraylab/brainsmash) on par-cellated left and right hemispheres. Parcellated surfaces, according to the 1000 cortical network parcellation described by Schaefer et al. (2018), are used instead of vertex-level data due to computational challenges of computing the required pairwise distance matrix for vertex-level data. As noted by Markello and Misic (2021), using such parcellations does not appear to compromise null model performance. For each hemisphere, a 501 × 501 geodesic distance matrix (which includes one parcel for the medial wall that does not need to be removed for BrainSMASH, unlike the spin test) is constructed using the SurfDist package in Python (Margulies et al. 2016). For BrainSMASH, we generate 1000 surrogate maps whose variograms are intended to be as close as possible to the empirical variogram of the observed cortical thickness maps. Variogram estimation includes pairwise distances in the bottom 25th percentile of possible pairwise distances, in accordance with Viladomat et al. (2014) and Burt et al. (2020). Since both tests of interest involve cortical thickness, for computational efficiency, we use the same 1000 surrogate cortical thickness maps tests against both sulcal depth and n-back in each age group.

#### 2.2.4 Testing correspondence within functional networks

Next, we test *H*_0(*spice*)_ and *H*_0(*b-smash*)_ and consider the same pairs of modalities as before in the left hemisphere, within canonical large-scale functional networks defined by Yeo et al. (2011) in their reported 7-network solution. Each network consists of a subset of the 1000 cortical parcellations described by Schaefer et al. (2018). For BrainSMASH, we increase the truncation threshold for pairwise distances included in variogram calculation to the 70th percentile, thus allowing variation between locations located farther apart to be incorporated in variogram calculation, following Burt et al. (2020)‘s methodology for assessment of spatial autocorrelation in subregions of the brain. To adjust for multiple comparisons across the seven age groups, we use a Bonferroni-adjusted *p*-value threshold for statistical significance (0.05/7 = 0.007). Because the main interest is in studying developmental changes throughout development in each network, we adjust for comparisons across age groups, but not across the Yeo networks nor the two tests.

## 3 Results

### 3.1 Type I error and power of the SPICE method

Simulation results are shown in Figure 4. Under both the null and alternative hypotheses, our proposed method performs well and as expected. The type I error (when 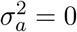) is close to 5% for all sample sizes, for all values of the vertex-level variance parameter, 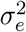, and for both settings involving *M*_2_(**v**) as either mean sulcal depth or mean n-back. Under the alternative hypothesis (when 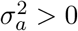), a higher signal-to-noise ratio (lower 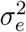, higher 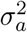) correspond to higher power, and power decreases incrementally as the signal-to-noise ratio decreases (higher 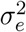, lower 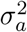). As expected, larger sample size and higher magnitude of population-level image similarity (Corr(*M*_1_(**v**), *M*_2_(**v**))) also improve statistical power.

**Figure 4:**
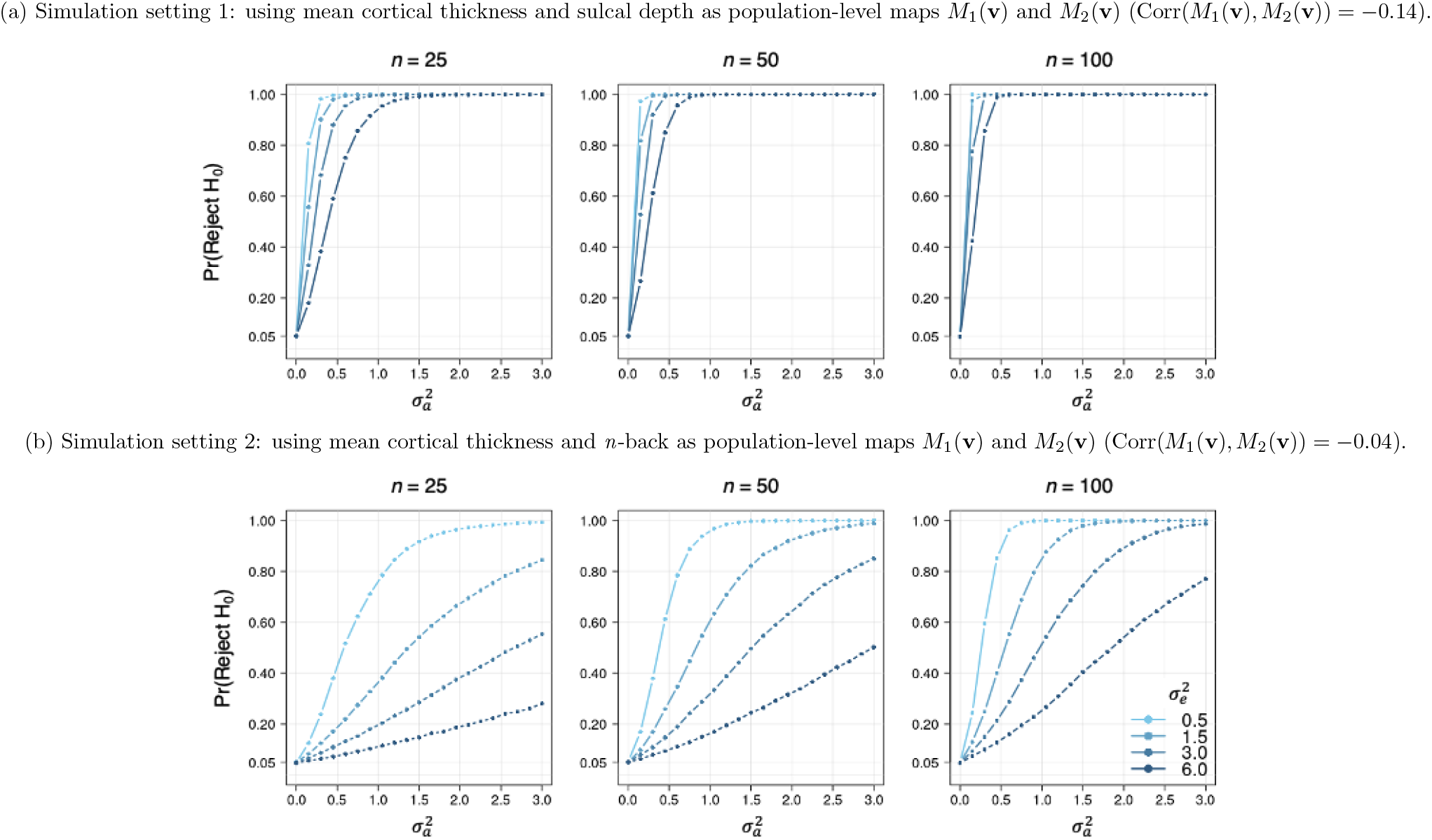
Power and type I error of the simple permutation-based intermodal correspondence (SPICE) test based on 5000 simulations. Each point is the rate of rejecting *H*_0_ (based on *p_spice_* < 0.05) from 5000 simulations of of the data (as shown in Figure 3) with unique combinations of parameters: sample size (*n* = 25, 50, or 100), subject-level variance (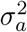, ranging from 0.0 to 3.0 in increments of 0.15), and vertex-level variance (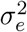, either 0.5, 1.5, 3.0, or 6.0). Both simulation settings involve using *M*_1_(**v**) as mean cortical thickness from a subset of 789 participants in the Philadelphia Neurodevelopmental Cohort. *M*_2_(**v**) is mean sulcal depth in setting 1 and mean *n*-back in setting 2.

### 3.2 Results and comparison of methods in the Philadelphia Neurodevelopmental Cohort

We first consider tests of intermodal correspondence for seven age groups in the left and right hemispheres. Unadjusted p-values are provided in Table 1. The SPICE test, BrainSMASH, and spin test all produce results that are consistent with what one would expect biologically: all three methods produce small p-value when testing sulcal depth versus cortical thickness (Table 1(a)), although BrainSMASH’s results in the right hemisphere for the youngest three age groups do not reach statistical significance based on our Bonferroni-adjusted *p*-value threshold of 0.003. None of the methods indicate significant correspondence between the *n*-back and cortical thickness maps (Table 1(b)). For BrainSMASH, we also verify that the surrogate maps’ variogram surrogate cortical thickness maps appear valid (Figure A1).

**Table 1:**
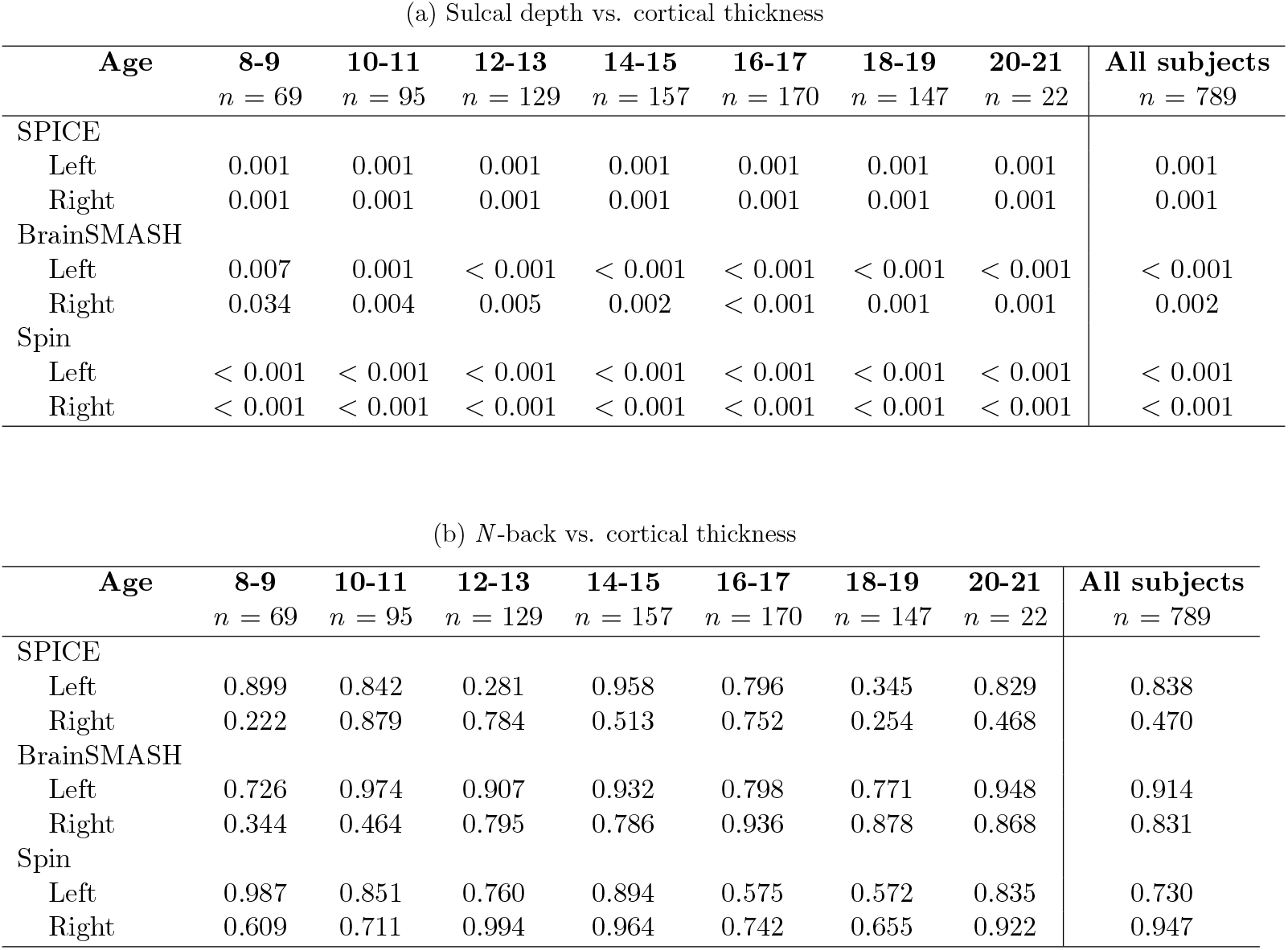
Unadjusted *p*-values from tests of intermodal correspondence in the left and right hemispheres using the SPICE, BrainSMASH, and spin methods. Null hypotheses for each method are summarized in Figure 1. Additionally, Figure A2 shows the null distributions and observed test statistics used to estimate each *p*-value below.

It is clear that intermodal coupling may occur with both anatomical and developmental specificity (Vandekar et al. 2016). For example, a positive result could be driven disproportionately by specific neuroanatomical sub-systems or specific age groups, suggesting the need for post-hoc regional or network analysis tests at higher anatomical or developmental resolutions. An important attribute of the SPICE test is its applicability for post-hoc testing in this sense. We therefore consider age-stratified tests of intermodal correspondence within functional networks from Yeo et al. (2011) with the SPICE test and BrainSMASH (plotted in blue and red, respectively, in Figure 5). As expected, the *p*-values for SPICE and BrainSMASH tests of correspondence for n-back versus cortical thickness (solid lines) appear uniformly distributed between 0 and 1, which we anticipate under their respective null hypotheses. However, we observe some discrepancies in their results for testing sulcal depth versus cortical thickness (dotted lines) within functional networks, despite the agreement of the two methods in tests conducted within full cortical hemispheres discussed before.

**Figure 5:**
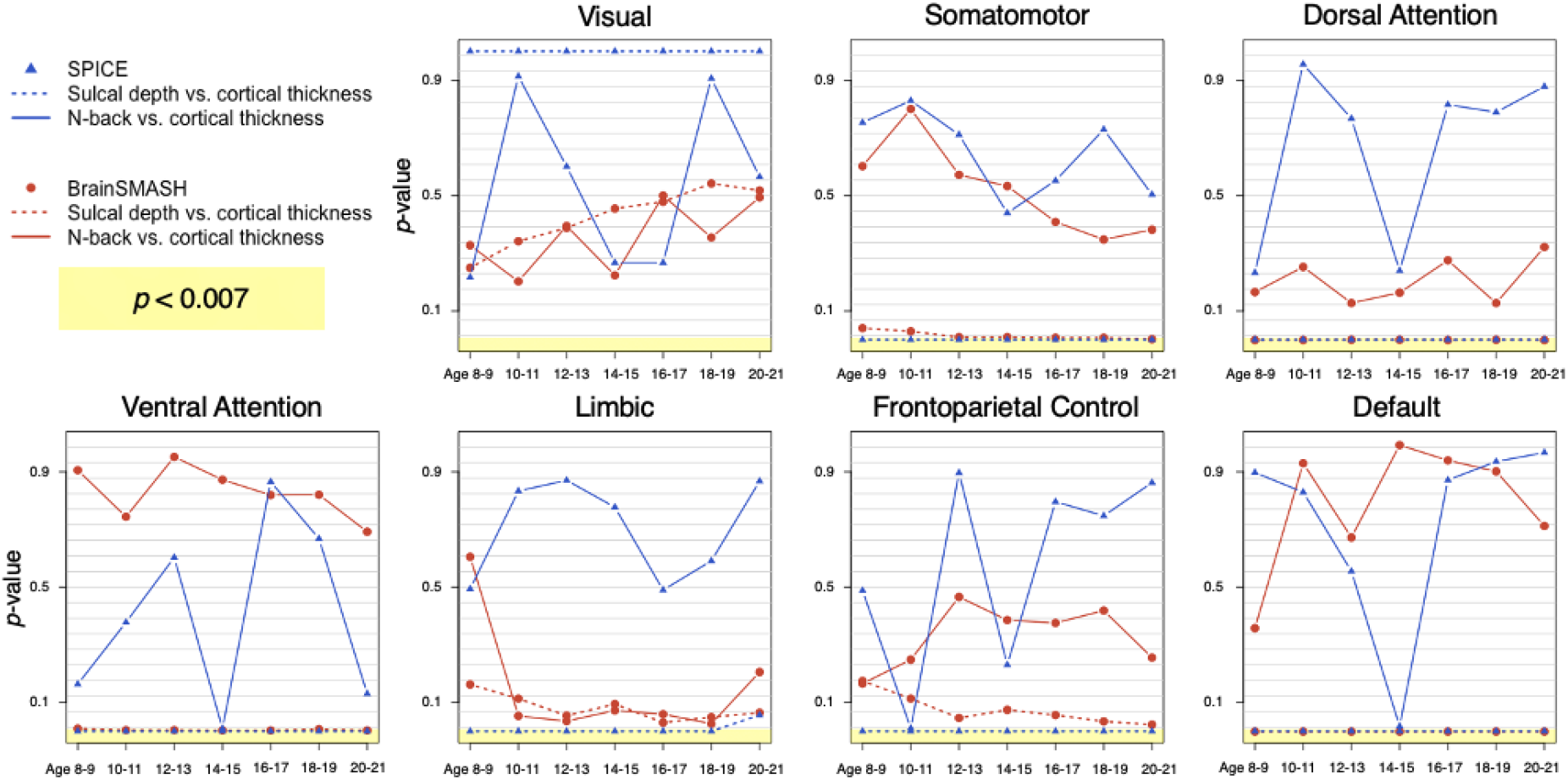
Unadjusted *p*-values from tests of intermodal correspondence within seven functional networks described by Yeo et al. 2011 in the left hemisphere for different age groups. We consider *p* < 0.007 to provide evidence against the null hypotheses (defined in Figure 1), after using a Bonferroni correction for comparisons across seven age groups.

Specifically, the SPICE test provides evidence against its null hypothesis (*H*_0(*spice*_)) of coupling between cortical thickness and sulcal depth in the somatomotor, dorsal attention, ventral attention, limbic, frontoparietal control, and default networks, with *p_*spice*_* < 0.007, the Bonferroni-adjusted threshold used for this analysis. BrainSMASH similarly provides evidence against its null hypothesis (*H*_0(*b-smash*)_) for coupling between sulcal depth and cortical thickness in the somatomotor (except ages 8-9 and 10-11), dorsal attention, ventral attention, and default networks, but not in the limbic and frontoparietal control networks.

The poor fit of the network-specific surrogate variograms, plotted in Figure A3, may give some insight for understanding these discrepancies. Specifically, we observe differences between the empirical and surrogate map variograms in all age groups for the dorsal attention, ventral attention, limbic, frontoparietal control, and default networks, suggesting that the surrogate cortical thickness maps in these networks failed to preserve the spatial autocorrelation structure of the target (i.e., original cortical thickness) map, undermining the results from tests in those networks, and suggesting that this method may not be reliable in the context of smaller brain regions.

It is also notable that neither the SPICE test nor BrainSMASH reject their respective null hypotheses for testing correspondence between sulcal depth and cortical thickness within the visual network, but that the non-significant p-values for the visual network are not uniformly distributed. This result is not surprising, given previous work from Vandekar et al. (2016), who found that in primary visual regions, we observe a complex relationship between these two modalities. These authors noted that the nonlinear nature of the association between sulcal depth and cortical thickness within the primary visual cortex violates the linearity assumption of their methodology for assessing intermodal coupling. It is plausible that this nonlinearity also explains SPICE’s and BrainSMASH’s failure to reject *H*_0(*spice*)_ and *H*_0(*b-smash*)_, respectively, within the visual network, as both methods currently use a linear measurement of similarity (Pearson correlation).

Lastly, although we did not hypothesize an association between the *n*-back task and cortical thickness, given the age-related regional heterogeneity in both cortical thickness growth curves (Tamnes et al. 2017) and also patterns of activation in the *n*-back task (Andre et al. 2016), differences in coupling across age groups and neuroanatomical systems are not entirely unexpected. Speculatively, the maturation of multi-modal association areas may be related to their differential activation in working memory tasks, which will be an interesting area for future work.

## 4 Discussion

In this paper, we introduce an intuitive and easy-to-implement method that leverages subject-level data to test and make inference about intermodal correspondence. Our method is complementary to those proposed by Alexander-Bloch et al. (2018), Burt et al. (2020), and other related methods described by Markello and Misic (2021), as the null hypotheses of these methods fundamentally differ (Figure 1) but may all provide meaningful insights about intermodal correspondence. While the spin test and BrainSMASH give a picture of population-level spatial alignment between two maps, our method considers subject-level intermodal correspondence, depending only on the plausible assumption that subjects are independent of one another. By approaching the correspondence problem from a new angle—emphasizing inference on subject-level associations as opposed to more general patterns of spatial alignment in group-averaged data—the SPICE test addresses five limitations of the earlier approaches.

Alexander-Bloch et al.’s spin test (i) can only be applied to surfaces with spherical representations (i.e., not volumetric data), (ii) does not specifically address the presence of the medial wall, which must be accounted for in practical implementations of the test, for example by disregarding cortical areas that overlap with the medial wall in spun maps when estimating the null distribution, and (iii) cannot be applied in small regions of the brain. While Burt et al.’s BrainSMASH can technically be applied in subregions of the brain, our findings suggest its generative null models may not be suitable when the small regions are irregularly shaped or disconnected, such as the widely used functional networks described by Yeo et al. (2011). One possible extension of BrainSMASH could incorporate nonparametric variogram estimation, as proposed by Ye et al. (2015), to account for both spatial proximity and functional connectivity in variogram estimation. Both the spin test and BrainSMASH are further limited in that they (iv) are only able to incorporate group-level data, precluding assessments of inter-subject heterogeneity (visualized in Figure 1). Finally, by conditioning on observed group-level maps, both of the previous methods would require (v) assuming covariance stationarity in order to take into account dataset-to-dataset variability or be used for generalizable statistical inference—that is, inference that formally considers the variability that results when sampling new data from the population. Without a treatment of this source of variability that does not rely on unrealistic assumptions, neither the spin test nor BrainSMASH may fully address concerns about the external validity of findings. The ability to directly address such concerns is a significant theoretical advantage offered by the SPICE test.

The SPICE test accounts for spatial autocorrelation to the extent that it is a feature of the underlying data-generating process. Spatially relevant information has already been encoded in the observed data, which renders moot any need for assumptions about stationarity. By conditioning on observed group-level data and altering its spatial structure in generating a reference null distribution, both the spin test and BrainSMASH necessitate explicitly modeling spatial autocorrelation or assuming covariance stationarity to make generalizable inference. In this sense, the SPICE test has the benefit of being readily usable for post-hoc analyses; if investigators decide to test correspondence between brain maps within subregions not considered in their primary analyses, the SPICE test would not require revisiting questionable assumptions about the spatial structure of the data when transitioning between testing more broadly or narrowly defined brain regions.

In addition to the SPICE test’s advantage of having minimal statistical assumptions, it is also clear that a method which allows for inter-individual variability would be advantageous in the neuroimaging field. Inter-individual variability in coupling between imaging-derived phenotypes has been shown to be sensitive to age, sex, and disease-related changes (Vandekar et al. 2016). However, we know of no systematic exploration of coupling across possible phenotype pairs, possibly due to the lack of a clear statistical framework for such a study. An example use case for the SPICE test is to prioritize phenotypic pairs for further analysis in large multimodal imaging studies, where coupling may be related to outcomes of interest such as psychopathology and individual differences (Karcher and Barch 2021). Given that inter-individual variability from both genetic and environmental sources is known to influence imaging phenotypes across modalities (Tooley et al. 2021; Fjell et al. 2015), higher within-subject coupling is consistent with “true” biological basis of intermodal coupling that also manifests marked inter-individual variability.

The SPICE test does not replace BrainSMASH, the spin test, or other methods that specifically wish to compare group-level surfaces, as SPICE is only applicable in settings where subject-level data are available. For example, the spin test and BrainSMASH may be used to test similarities between different atlases (e.g., Yeo versus Desikan), and the SPICE test would not be applicable in such studies.

In future work, we hope to consider more comprehensive testing strategies, including the use of different correspondence metrics, since the choice of statistic should be appropriate for the modalities under consideration. For example, while the Pearson correlation has known limitations when used in sparse data such as connectivity matrices, a geodesic distance measure between two connectivity matrices may be more suitable (Venkatesh et al. 2020). In addition, since the null hypothesis for the SPICE test is framed in terms of equality in distributions of a correspondence measure (1), one may consider additional test statistics (for example, the Kolmogrov-Smirnov statistic) so that the test can explicitly consider the full distribution function, rather than focusing on the mean. Implementing the SPICE test using a Pearson correlation (or other average measurement) tests only one aspect of the distribution—testing a sufficient but not necessary condition for equivalence in distributions. A rank-based measure of similarity (e.g., the Spearman correlation coefficient) would be another option more sensitive to nonlinear associations.

Finally, since our method involves permuting subject-level maps in generating a null distribution, in future work we plan to consider implications of subject exchangeability on inference using our method. Various approaches to preserve exchangeability (e.g., defining blocks within which subjects may be considered “exchangeable”) may be adapted from methods described by Winkler et al. (2014).

## 5 Conclusions

The SPICE, BrainSMASH, and spin methods may all support similar findings despite their different null hypotheses. However, the SPICE test is the most flexible when it comes to analyzing subregions of the brain, particularly when assumptions regarding the structure of spatial autocorrelation can pose obstacles to generating surrogate maps intended to preserve those complex structures and it is the only method that is able to consider subject-level data. Depending on the question of interest and the available data, using a combination of these three methods may be beneficial in future neuroimaging experiments to obtain a more complete picture of intermodal correspondence.

## Data availability statement

Neuroimaging data were acquired as part of the Philadelphia Neurodevelopmental Cohort (PNC), a research initiative at the University of Pennsylvania Brain and Behavior Laboratory and Children’s Hospital of Philadelphia. PNC data are publicly available in raw format at https://www.ncbi.nlm.nih.gov/projects/gap/cgi-bin/study.cgi?study_id=phs000607.v3.p2. Additionally, R code for reproducing our simulations can be found at https://github.com/smweinst/spice_test.

## Funding statement

This work was supported by the following NIH grants: R01MH123563 (S.N.V.), R01MH107235 (R.C.G.), R01MH119219 (R.C.G. and R.E.G.), R01MH113550 (T.D.S.), R01EB022573 (T.D.S.), R01MH120482 (T.D.S.), RF1MH116920 (T.D.S.), K08MH120564 (A.F.A.B.), R01MH112847 (R.T.S. and T.D.S.), and R01NS060910 (R.T.S.). This study was supported by the intramural research program of the National Institute of Mental Health (NIMH) (project funding: 1ZIAMH002949) (A.R.). This work was also supported by the National Science Foundation Graduate Research Fellowship Program (S.M.W.).

## Conflict of interest disclosure

The authors declare no competing interests.

## Ethics approval statement

The institutional review boards of the Hospital of the University of Pennsylvania and Children’s Hospital of Philadelphia approved protocols for the PNC.

## Patient consent statement

Participants and/or their parents provided informed consent to be included in the PNC.

## Permission to reproduce material from other sources

N/A

## Clinical trial registration

N/A

## Author contributions

**S.M.W.:** conceptualization, data analysis, software, and writing. **S.N.V.:** conceptualization, writing/review of manuscript. **A.A.:** data processing and review/editing of manuscript. **T.M.T.:** data processing and review/editing of manuscript. **T.R.F.:** data analysis, software, and review/editing of manuscript. **R.C.G.**: data collection, data processing, and review/editing of manuscript. **R.E.G.**: data collection, data processing, and review/editing of manuscript. **A.R.**: conceptualization and review/editing of manuscript. **T.D.S.**: conceptualization, data collection, data processing, and review/editing of manuscript. **A.F.A.B.:** conceptualization, data analysis, writing/review of manuscript, and supervision. **R.T.S.:** conceptualization, data analysis, writing/review of manuscript, and supervision.

## Appendix

### A1 Imaging protocols

#### A1.1 Acquisition and pre-processing of structural imaging data

All subjects underwent magnetic resonance imaging (MRI) in the same Siemens TIM Trio 3 tesla scanner with a 32-channel head coil and the same imaging sequences and parameters, described in detail by Satterthwaite et al. (2014) and Satterthwaite et al. (2016). For structural imaging, the protocol included a magnetization-prepared, rapid-acquisition gradient echo (MPRAGE) T1-weighted structural image with a voxel resolution of 0.9 × 0.9 × 1 mm. To ensure adequate quality of images included in our analysis, quality assurance was independently rated by three experienced image analysts, as described by Rosen et al. (2017).

Cortical reconstruction of T1-weighted structural images was implemented using FreeSurfer (version 5.3), which included template registration, intensity normalization, and inflation of cortical surfaces to a template. Cortical measurements were resampled to the fsaverage5 atlas, consisting of exactly *V* = 10,242 vertices in each hemisphere for every subject. We quantified cortical thickness as the minimum distance between pial and white matter surfaces (Dale et al. 1999) and sulcal depth as the height of gyri (with positive and negative values indicating outward and inward movement, respectively) (Fischl et al. 1999).

#### A1.2 Acquisition and pre-processing of *n*-back task sequence

Protocols for functional MRI (fMRI) included task-based BOLD scans using a single-shot, interleaved multi-slice, gradient-echo, echo planar imaging sequence, with a voxel resolution of 3 × 3 × 3 mm voxels and 46 slices. Preprocessing was implemented using the eXtensible Connectivity Pipeline (XCP) Engine (Ciric et al. 2018). In the current study, we consider the fractal *n*-back task sequence, which involves presenting subjects with a series of stimuli (geometric pictures on a screen) with instructions to press a button if the current stimulus matches the *n*th previous one. For example, in a 1-back sequence, participants would be instructed to recognize whether a current stimulus matches the one that appeared last; a 2-back sequence would involve recognizing whether the current stimulus matches the one that appeared two stimuli ago. A 0-back sequence is also considered, in which subjects are instructed to simply press a button each time a stimulus appears (regardless of whether it matches a previous stimulus). In our present analysis we consider maps that represent the percentage change in activation between the 2-back and 0-back sequences, which has been previously shown to isolate executive network function (Ragland et al. 2002).

We used the FEAT software tool in the FSL library to fit first-level general linear models on the *n*-back time series data (Jenkinson et al. 2012). Three first level models for each subject considered contrasts to assess change in working memory load between 1-back and 0-back, 2-back and 1-back, and 2-back and 0-back. Activation maps representing the percent change between the 2-back and 0-back sequences were projected into the fsaverage5 template with exactly 10,242 vertices per hemisphere per subject.

#### A1.3 Use of cortical and network parcellations from Yeo et al. (2011) and Schaefer et al. (2018)

Images converted to the fsaverage5 template consisted of exactly 10,242 vertices per hemisphere. We then identified the vertices belonging to each of the 1000 parcellations suggested by Schaefer et al. (2018) as well as each of the seven networks proposed by Yeo et al. (2011), available for download from GitHub (https://github.com/ThomasYeoLab/CBIG/tree/master/stable_projects/brain_parcellation/Schaefer2018_LocalGlobal/Parcellations/FreeSurfer5.3/fsaverage5/label).

### A2 Supplemental figures

**Figure A1:**
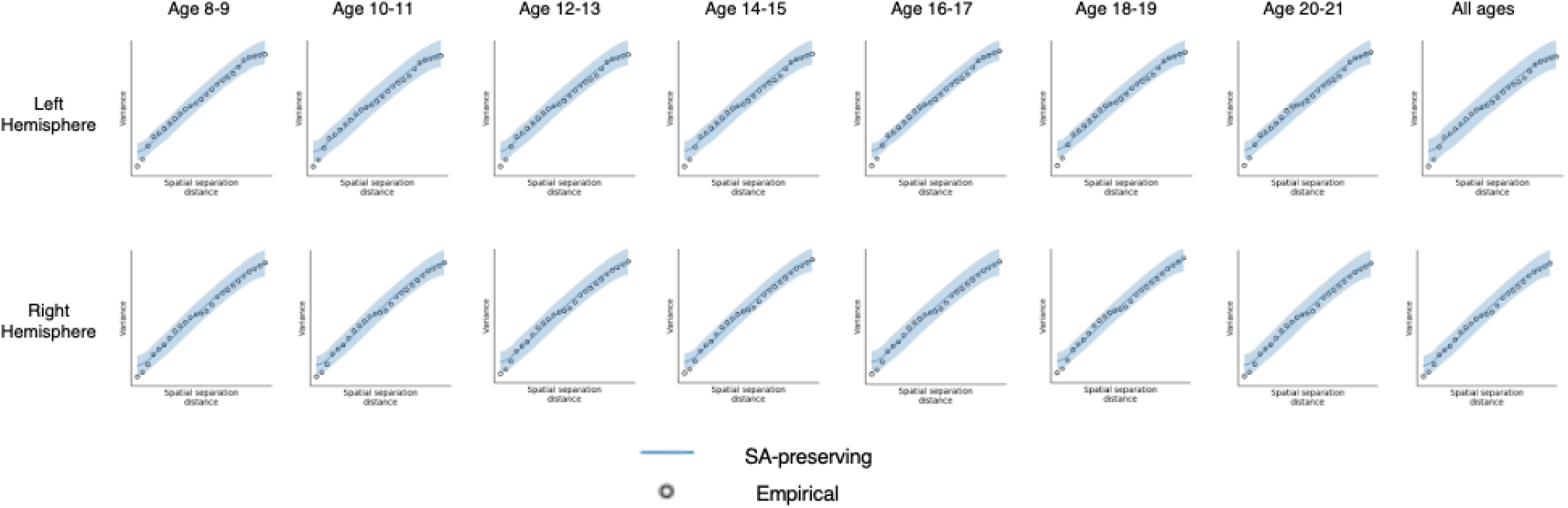
Empirical and surrogate (no. surrogates = 1000) variograms of parcellated cortical thickness measurements. These variograms are constructed to assess the reliability of the Brain Surrogate Maps with Autocorrelated Spatial Heterogeneity (BrainSMASH) method in testing for intermodal correspondence between cortical thickness and sulcal depth and cortical thickness and n-back in the left and right hemispheres (parcellations from Schaefer et al. (2018)). The horizontal axis of each figure indicates the spatial separation distance (*d*), and the vertical axis describes the variation between the cortical thickness measurements observed in parcels separated by distance *d*.

**Figure A2:**
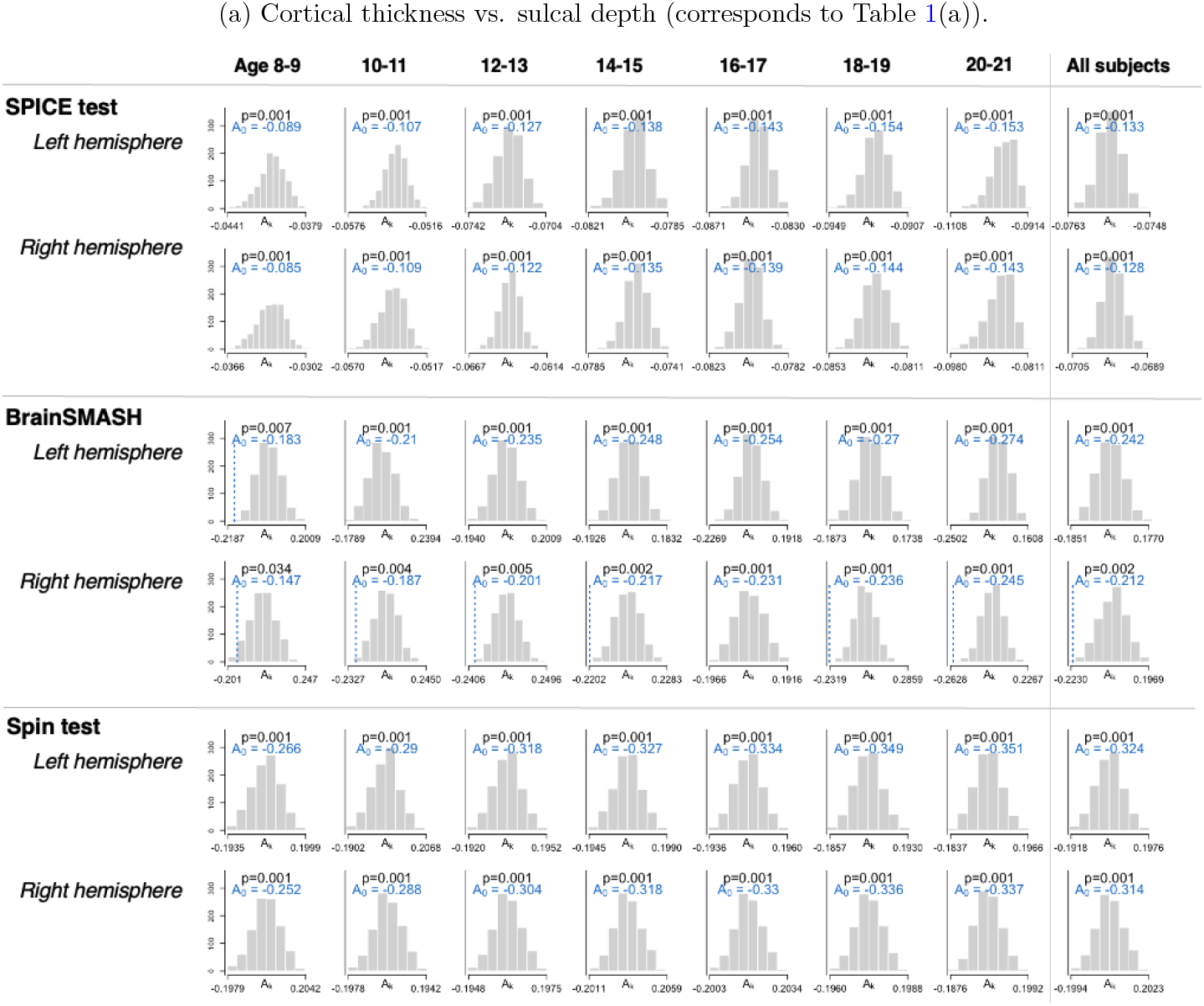
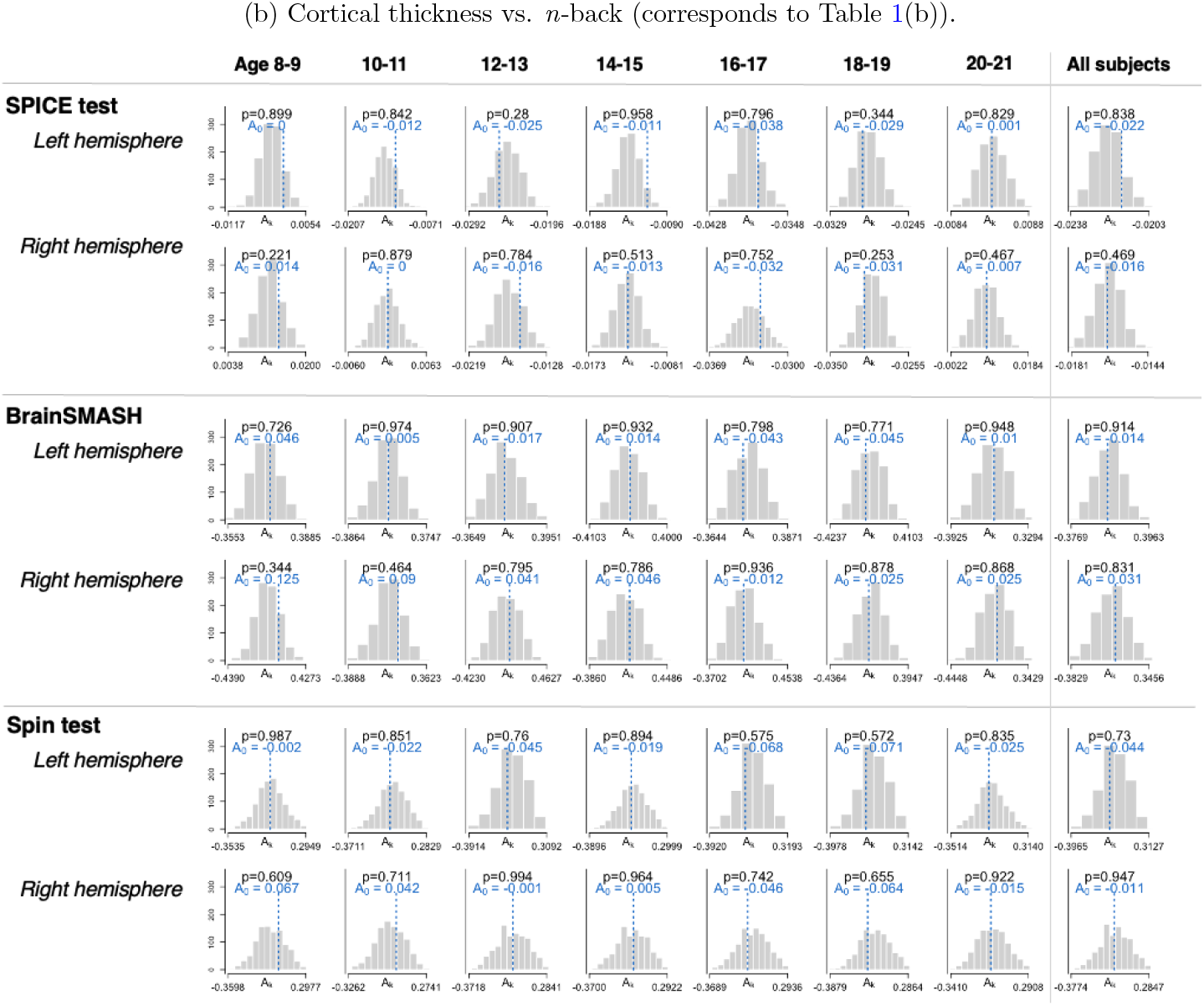
Null test statistic distributions corresponding to results shown in Table 1 for the SPICE test, BrainSMASH, and spin test. The observed test statistic, *A*_0_, is plotted or written in blue. (*A*_0_ is not plotted when it falls outside the range of the null test statistics.) Note: the observed test statistics for BrainSMASH and the spin test are not identical, even though both these methods use group-level surfaces, since the spin test removes the medial wall before calculating intermodal correspondence.

**Figure A3:**
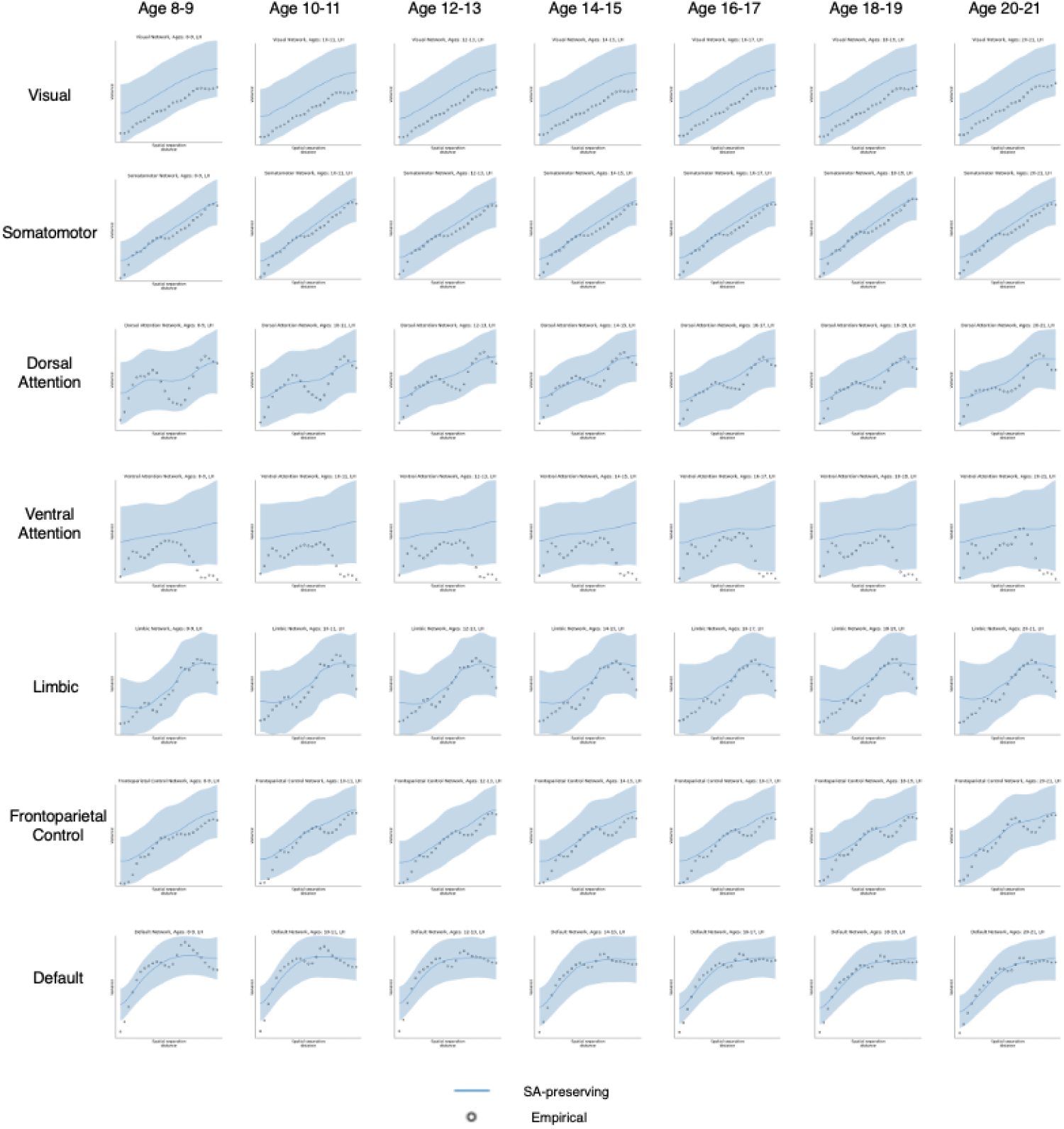
Empirical and surrogate (no. surrogates = 1000) variograms of parcellated cortical thickness measurements within seven functional networks, according to Yeo et al. (2011). These variograms are constructed to assess the reliability of BrainSMASH in testing for intermodal correspondence between cortical thickness and sulcal depth and cortical thickness and *n*-back within age-stratified groups. The horizontal axis of each plot indicates the spatial separation distance (*d*); the vertical axis describes the variation between cortical thickness measurements observed in parcels separated by the distance *d*.

## Notes

https://github.com/smweinst/spice_test

### Competing Interest Statement

The authors have declared no competing interest.

